# Formal string instrument training in a class setting enhances cognitive and sensorimotor development of primary school children

**DOI:** 10.1101/829077

**Authors:** Clara E. James, Sascha Zuber, Elise Dupuis-Lozeron, Laura Abdili, Diane Gervaise, Matthias Kliegel

## Abstract

This randomized controlled trial shows for the first time that focused musical instrumental practice as compared to traditional sensitization to music provokes robust cognitive and sensorimotor transfer effects. Over the last two years of primary school (10-12-year-old children), sixty-nine children received biweekly musical instruction by professional musicians within the regular school curriculum. The intervention group learned to play string instruments, whereas the control group, peers in parallel classes, was sensitized to music via listening, theory, and some practice. Broad benefits manifested in the intervention group as compared to the control group for working memory, attention, processing speed, cognitive flexibility, matrix reasoning, sensorimotor hand function and bimanual coordination Apparently, learning to play a complex instrument in a dynamic group setting impacts development much stronger than classical sensitization to music. Our results therefore highlight the added value of intensive musical instrumental training in a group setting, encouraging general implementation in public primary schools, better preparing children for secondary school and for daily living activities.

**Highlights:**

- Music practice produces multiple cognitive transfer effects in school children
- Making music causes stronger cognitive benefits than sensitization to music
- Music practice in a class setting improves executive functions and abstract thinking
- Playing string instruments enhances fine manual dexterity and bimanual coordination

## 1. Introduction

Practicing a complex instrument regularly and actively over extended periods of time may provoke positive transfer effects on basic and higher order cognition as well as on sensorimotor skill in children (Bergman Nutley, Darki, & Klingberg, 2014; Costa-Giomi, 2004; Martins, Neves, Rodrigues, Vasconcelos, & Castro, 2018; Moreno et al., 2011; Roden, Kreutz, & Bongard, 2012; Schellenberg, 2004, 2006; Schlaug, Norton, Overy, & Winner, 2005; Tierney, Krizman, & Kraus, 2015). Some studies even suggest long-term effects of musical practice during childhood (Balbag, Pedersen, & Gatz, 2014; Hanna-Pladdy & MacKay, 2011; Moreno, Lee, Janus, & Bialystok, 2015; Schellenberg, 2006; White-Schwoch, Woodruff Carr, Anderson, Strait, & Kraus, 2013). Yet, importantly, as summarized by Dumont and colleagues’ (Dumont, Syurina, Feron, & van Hooren, 2017) recent review, the available literature is largely inconclusive because of the lack of genuine randomized controlled trials and active control groups. The authors also observed a great heterogeneity across the different studies, concerning group size, intensity and nature of the music regimens. Of the 46 studies on music interventions that the Dumont review comprised, only two used an RCT design. Neither of these two studies, lasting 6 months and 6 weeks respectively, could show cognitive benefits in the music groups as compared to art classes (Flaugnacco et al., 2015; Mehr, Schachner, Katz, & Spelke, 2013).

Specifically concerning group training, Rickard, Bambrick, & Gill (2012) investigated school-based instruction in young adolescents over 5-6 months in a pseudo-randomized study. No convincing developmental benefits of music lessons in a class setting manifested, compared to control groups that received drama and art classes. The proposed music trainings (Rickard et al., 2012) involved conscious listening and introduction to basic musical concepts, playing and improvising on different instruments and learning different musical notations. Another group setting study (Degé, Wehrum, Stark, & Schwarzer, 2011) compared children in similar age groups as in the current study (9-11 years), after two years of intensive school music training, with a passive control group (Degé et al., 2011). Participants were not assigned randomly to the music groups. The authors found evidence for enhanced short term auditory and visual memory in the music groups. Their musical regimen did involve playing an instrument. Slater and colleagues (2014), in a pseudo randomized group setting, observed a catch-up in reading performance in low-income children after one year of music training comprising focused instrumental training as well as musical theoretical education. However, they used a passive control group and only children that desired to participate were included in the music group, inducing a motivational bias.

Jaschke, Honing and Scherder (2018a) observed in a cross-sectional study in 6 year old children that extra-curricular exposure to a musically enriched environment (listening) did not provoke significant relationships with cognitive function, although a trend with verbal intelligence appeared.

From all these studies, we conclude that it seems predominantly focused instrumental practice, i.e. learning to play a complex musical instrument over an extended period of time that constitutes the main driving force for far transfer to basic and higher order cognitive processing and sensorimotor skills. Yet, so far, this has never been investigated systematically within a long-term RCT with an active control group, in an intra-curricular thus natural class setting, involving children of different backgrounds, using a large behavioral battery covering distinct developmental domains.

Available evidence of beneficial musical practice effects on cognitive child development predominantly concerns children of parents with a high socioeconomic and educational background (Corrigall & Schellenberg, 2015) and typically results from private lessons. Additionally, most of the time, the child is interested to learn a musical instrument, inducing a motivational bias (Corrigall, Schellenberg, & Misura, 2013). Evaluation of beneficial transfer effects restrains in general to a limited number of capacities or skills. and randomized controlled trials (RCTs) with active control groups are scarce.

Here, we compared children who intensively practiced different string instruments in a class setting within a specific Orchestra in Class (OC) program, to peers in parallel classes that received the same amount of musical instruction, also within an entire class, but lacking focused training on a complex musical instrument. Entire existing classes were assigned randomly to the OC and the Control programs. The study took place in public primary schools in popular neighborhoods in the Geneva area, avoiding confounding music effects with effects of socioeconomic background.

We anticipated that cognitive functions strongly involved in musical practice like working memory, attention, information processing, cognitive flexibility and abstract reasoning, as well as fine sensorimotor function would provoke enhanced positive transfer effects in the OC group as compared to the control group.

## 2. Materials and Methods

### 2.1 Participants

Sixty-nine primary schoolchildren participated in the study (M at baseline = 10.18 years; SD = 0.31; 41 girls). We recruited the children at the establishment were the OC program was integrated in the regular curriculum in French-speaking Switzerland, and in neighboring public schools, in a popular neighborhood, therefore hosting children of varying ethnic and socio-economic backgrounds. At baseline, before the interventions, the children almost finished their sixth year of elementary school, one of eight consecutive years covering ages between four and twelve years approximately. We excluded any children who followed regular or protocolled music practice outside the school curriculum before the study. Seven children were left-handed, three in the control group and four in the OC group (see supplementary Table 1, at the end of the manuscript). We integrated handedness in the linear mixed models, controlling for this factor (see 2.9.1 Linear Models).

The children and their parents or caregivers signed an informed consent. The ethics commission of the Faculty of Psychology and Educational Sciences of the University of Geneva approved the protocol, in agreement with the ethical standards of the Helsinki declaration.

### 2.2 Musical interventions

Assignment of whole classes to the OC group or Control group was random (cluster randomization; Mazor et al., 2007). The most important advantage of the cluster randomization in our context is that the intervention is tested under representative natural conditions (Mazor et al., 2007). If the Orchestra in Class program would be generalized to more primary schools, the interventions would take place in existing classes. Additionally, we controlled for baseline differences in the Linear mixed models (see 2.9.1 Linear Models). Finally, we excluded all children that followed music lessons prior to the interventions from the study. The intervention group comprised 34 children (two classes), the control group 35 (two classes).

#### 2.2.1 OC group

Children received OC courses within a whole class two times per week during 45 minutes within their own school during the last two years of primary school. First the children were assigned their instrument (violin, viola, cello or double bass). At the beginning playing involved bowing open strings smoothly “legato” (without using the fingers of the left hand) and using pizzicato (plucking the strings with the right hand), in order to familiarize the child with the instrument. Then progressively the fingers of the left hand were used to stop the strings, first while playing pizzicato, in order to concentrate on the fingers of the left hand, and later on in combination with more and more diversely articulated use of the bow. Meanwhile rhythms evolved from very simple regular to more complicated and irregular ones. After three months the children could take their instruments home and were encouraged to practice on a voluntary basis. After one year the average child could play on all strings with all four fingers of the left hand (the thumb stabilizes the neck of the instrument) and could use varied bowings. Score reading was gently initiated during the first year, but really applied in the second year. At all times, from the very beginning, ensemble playing remained a priority, the child was made aware constantly of being part of a whole within a polyphony.

Learning followed three paths, by imitating the teacher and later on also by reading the score. Finally, emulation among students was also an important vehicle of learning.

Small concerts and events stimulated the children and gave purpose to their learning, including two study weekends with a final concert in front of the families.

The OC teachers were professional string players, specifically prepared for this way of teaching at the Accademia d’Archi, an accredited music school in Geneva., specialized in string instrument education.

#### 2.2.2 Sensitization to music

In the Geneva Canton all children in this age group attaining public schools, receive two times 45 minutes of musical education per week, and so did our control group during the last two years of primary school. The education is best described as “sensitization to music” and involves listening actively, learning some theory, singing together and playing small percussive instruments or the recorder. The teachers were professional musicians that received training to provide musical education in a school setting. The children, like the OC group, also participated in class performances for the parents.

### 2.3 Procedure

Master students of the Psychology Department of the Geneva University tested all children individually, within the school that the child attended. Prior to testing, the experimenters informed the children that they would participate in a study of the University of Geneva on child development. The experimenters encouraged each child to ask questions and emphasized that he/she may ask for a break at any time. All experimenters were well trained (two 3-hour sessions) to pass the tests correctly and uniformly beforehand.

The tests were administered in pseudo-randomized order, in a time window of approximately two hours, separated by breaks. Total testing time took one hour and 30 minutes, separated by breaks. The children received a small gift at the end of each session.

### 2.4 Materials

All types of tests, measured variables, acronyms of the tests and involved abilities are resumed in Table 1.

**Table 1.**
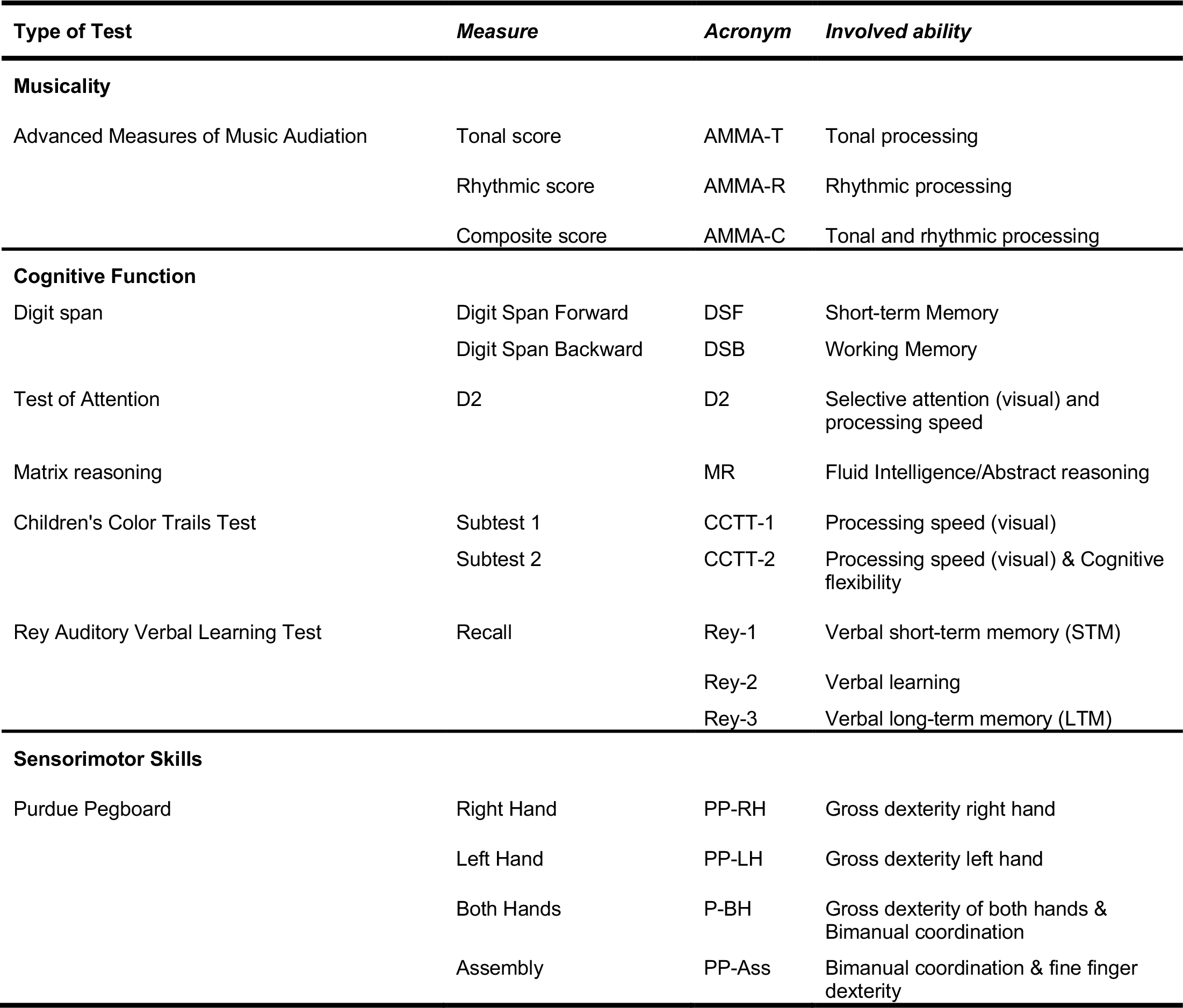
Types of tests, Measures, Acronyms and involved abilities of all items of the test battery.

#### 2.4.1 Music Audiation

As just possessing good discriminatory skills for pitch and rhythm is not sufficient to evaluate musical aptitude, we administered the “Advanced Measures of Music Audiation” (AMMA; Gordon, 1989). This test requires the capacity to group individual notes into “Gestalts” and to form expectancies thus evaluating “auditory structuring” (Karma, 2007). The AMMA test does not require any prior musical knowledge or skills and is suited to evaluate musical aptitude in pre-adolescents up to professional musicians (Grades 7 to Adult). The test encompasses 30 trials consisting of pairs of musical melodies presented over headphones via the computer. The children judged for each pair whether the melodies were identical or different and if they considered that the two melodies of the pair were different, they had to indicate whether the difference was melodic or rhythmic. Tonal and rhythmic differences never occurred together. Among the 30 pairs, ten pairs are identical, ten melodically different and ten rhythmically different. The first phrase always contains exactly the same number of notes as the second. Scoring is divided in a tonal sub-score, a rhythmic sub-score and a composite score that is a combination of both tonal and rhythmic scores. The performance on this test represents an interaction between innate musical potential and exposure to musical environments. Because the scoring system penalizes errors, we advised the children not to respond if they were not sure. The answer sheet that was filled in by the experimenters to prevent errors, thus contained four columns: identical – melodic difference – rhythmic difference – I don’t know. After explaining the concepts of melody and rhythm in a plastic way, the children passed three training trials that were discussed with them to ensure that they understood the instructions. We computed a tonal a rhythmic and a composite standard score, the latter composed of both tonal and rhythmic scores, according to the AMMA manual (Gordon, 1989), thus applying a subtraction of points for wrong answers. From these standard scores we inferred percentile rank scores (category High School Students) according to the AMMA manual (Gordon, 1989), that we used for the analyses.

#### 2.4.2 Digit span forward and backward

All children passed the “digit span” subtest of the Wechsler Intelligence Scale for Children - Revised (WISC-R; (Wechsler, 2005)). In this test the participant listens to series of digits with increasing length and has to repeat them orally. In the Digit Span Forward task (DSF) in direct order, in Digit Span Backward (DSB) in reverse order. To ensure a regular time course (1 sec. per digit) and identical pronunciation of the presented material for all participants, we pre-recorded the spoken series. DSF and DSB assess distinct but interdependent cognitive functions (Grégoire, 2009). DSF evaluates essentially short-term auditory memory, whereas DSB evaluates principally the ability to manipulate verbal information while temporarily stored, thus auditory working memory capacity. Two series of digits (one for each task), progressively increasing in length, thus in difficulty, were presented. The children first performed the DSF (span size from 2 up to 9) then the DSB task (span size from 2 up to 8). The task was interrupted if the child made two successive mistakes with the same number of digits, i.e. at the same level of difficulty. Each correct answer counted for one point.

#### 2.4.3 D2 test of Attention

To assess children’s selective visual attention, sustained attention and visual scanning speed (processing speed) we administered the D2 Test of Attention (Brickenkamp & Zillmer, 1998). Stimuli consisted of letters d or p accompanied by one or two apostrophes above and/or below the letter, presented on a paper sheet with 14 rows of 47 stimuli. The participant should cross-out all the d’s accompanied by exactly two apostrophes (i.e. two apostrophes above, two apostrophes below or one above and one bellow the d), without crossing-out any of the distractors (d’s accompanied by only one apostrophe and all p’s). To familiarize the children with the task, they first performed a practice row of 22 trials. For the actual task, children started working on the first row and were asked to switch to the next row every 20 seconds. The outcome measure (D2) used provides the total number of correctly marked items minus the number of errors and omissions.

#### 2.4.4 Matrix reasoning

In order to appraise abstract reasoning, we applied the Matrices subtest of the WISC-IV (Wechsler, 2003). Since musical phrases develop over time according to musical grammar, like language (Patel, 2012), we consider that some abstract thinking is involved in producing and processing music (Jaschke, Honing, & Scherder, 2018b). The test consists of different sheets with series of three images (e.g. three oval shapes). The child should detect the image that correctly completes the series (e.g. another oval shape) among four distractors (e.g. other shapes). Prior to the task, the children went through three practice trials, to assure they understood the instructions. For the real task, the sheets gradually increased in difficulty. The task was interrupted when the child answered four out of five consecutive sheets incorrectly. The number of correctly answered sheets constitutes the final score.

#### 2.4.5 Children’s Color Trails Test

The Children’s Color Trails Test (CCTT; Llorente, 2003) consists of two subtests. In the CCTT-1 test children are given a sheet on with 15 circles containing the digits “1” to “15” are presented. The child should connect the digits in increasing order with a pencil as rapidly and correctly as possible. All circles with even digits were colored yellow, whereas the circles with odd numbers were colored pink. For the CCTT-2 test, children performed the same task with the difference that for each digit (except for number “1”) two circles were presented on the sheet, one colored yellow, the other colored pink. Children were instructed that in addition to connecting the digits in increasing order, the colors of the circles would have to alternate for each digit (the pink “1” had to be connected with the yellow “2”, which had to be connected to the pink “3” etc.). The CCTT-1 evaluates simple visual processing speed, the CCTT-2 visual processing speed plus cognitive flexibility. Both subtests started with an 8-digit practice sheet for familiarization purposes. We computed standard scores as outcomes for both subtests (M = 100; SD = 15). For the analyses we used associated percentile scores.

#### 2.4.6 Rey Auditory Verbal Learning Test

A list of 15 unrelated words was presented orally to the children five times in a row (Bean, 2011; Rey, 1964). Each time, the children should repeat as many correct words out of the list as possible after a short break of approx.10 seconds. The time limit for recollection was set at one minute for the first trial, and for trials 2-5 to one minute and 30 seconds. Each time the list was read aloud first. Trials 2-5 were performed immediately after trial 1. After a delay of about 50 minutes, the child should again cite as many words as possible from the list, but this time, without their oral presentation before. The scores represent each time the number of correctly repeated words. We composed the following measures: 1) the score of trial 1 (Rey-1) evaluating verbal short-term memory (STM), 2) the mean score of trials 2-5 (Rey-2), evaluating verbal learning and 3) a score of delayed recall (Rey-3), evaluating verbal long-term memory (LTM). At each time point different lists were used.

#### 2.4.7 Purdue Pegboard

The Purdue Pegboard test administered according to the Lafayette manual (Lafayette, 1999) serves to measure manual gross and fine dexterity as well as bimanual coordination. The Pegboard contains 2 parallel rows of 25 vertically oriented holes. Two cups on top of the board contain pegs (diameter one mm), collars and washers. After a familiarization phase, the children inserted as many pegs as possible in the holes in 30 seconds, from top to bottom, first with their right hand (PP-RH), then with their left hand (PP-LH) and finally with both hands simultaneously (PP-BH). PP-RH and PP-LH evaluate gross hand dexterity, PP-BH gross hand dexterity plus bimanual coordination. Then, again after a familiarization phase, the children performed an assembly task working with both hands together, placing as many assemblies in the holes as possible in one minute (PP-Ass). This sub-task requires bimanual coordination in combination with fine finger dexterity. One assembly consisted of a peg, a collar and two washers (four elements) to be placed into one hole in a specific order. Four scores were collected, corresponding to the number of pegs placed (PP-RH, PP-LH, PP-BH) and the number of correctly inserted elements placed during the PP-Ass task.

### 2.8 Missing Data

We report, depending on the test (see supplementary Table 1 for more details) one missing value at most at T0 (AMMA and Purdue tests), 2 missing values at T1 for all tests and from 5 to 6 missing values (AMMA test) at T2.

### 2.9 Analyses

#### 2.9.1 Linear mixed models

Linear Mixed models are a generalization of ANOVA and linear regression for situations where measures are repeated on the same individuals. They are flexible and powerful statistical models that can take into account several levels of clustering, continuous and qualitative explanatory variables as well as unbalanced data (Bates, Mächler, Bolker, & Walker, 2015).

Three linear mixed model equations were composed for each test in order to model the evolution of scores over time (T1 vs T2) for each child from each group, with a random intercept for each child. All models comprised score at T0, age at T0, gender and handedness in order to control for these factors.

Model 1: Time * Group Interaction, to verify whether Groups evolve differently over Time: *Score ~ T0 + SEX + Lat + Age_2016 + Time + Group + Time:Group*

Model 2: Effect of Time & Group: *Score ~ T0 + SEX + Lat + Age_2016 + Time + Group*

Model 3: Effect of Time: *Score ~ T0 + SEX + Lat + Age_2016 + Time*

The lme4 and the emmeans package of the software R (3.6.0) served to estimate the model parameters (https://CRAN.R-project.org/package=emmeans), freely available at http://www.R-project.org (Bates et al., 2015).

#### 2.9.2 Likelihood-ratio tests

The statistical significance of the Interaction effect Time * Group and of the factor Group was assessed using likelihood-ratio tests (Pinheiro & Bates, 2006) To do so, models with and without the factor of interest (Interaction Time * Group; Group) were compared. So, we tested the significance of the interaction Time * Group by comparing the 1st and the 2nd model with a likelihood-ratio test. In the same way we tested the effect of the group by comparing the 2nd and the 3rd model.

This procedure resulted in values of the observed chi-square test statistic, associated p-values, and effect sizes (partial Rsquare/r2 at the level of the test and Rsquare/r2 at the level of the model) for Interaction Group * Time and Group (see Table 2).

**Table 2.** Results of the two likelihood tests comparing the linear mixed models, expressed by means of observed chi-square test statistics, associated p-value, and effect size (partial Rsquare/r2 at the level of the test and the model). The effect size is only provided in the case of a significant effect.

| Measure | Likelihood test | Chi_2 | Df | p-value | Partial R<sup>2</sup> | R<sup>2</sup>_model |
| --- | --- | --- | --- | --- | --- | --- |
| AMMA-T | Interaction Time*Group | 0.406 | 1 | 0.524 |  | 12.5 |
|  | Group | 5.028 | 1 | **0.025** | 9.4 |  |
| AMMA-R | Interaction Time*Group | 0.024 | 1 | 0.877 |  | 21.7 |
|  | Group | 8.013 | 1 | **0.005** | 13.1 |  |
| AMMA-C | Interaction Time*Group | 0.167 | 1 | 0.683 |  | 22.4 |
|  | Group | 10.368 | 1 | **0.001** | 15.8 |  |
| DSF | Interaction Time*Group | 0.001 | 1 | 0.978 |  | 51.1 |
|  | Group | 1.067 | 1 | 0.302 | 1.6 |  |
| DSB | Interaction Time*Group | 3.225 | 1 | 0.073 | 4.8 | 39.8 |
|  | Group | 4.650 | 1 | **0.031** | 7.0 |  |
| D2 | Interaction Time*Group | 1.392 | 1 | 0.238 |  | 64.8 |
|  | Group | 4.044 | 1 | **0.044** | 5.9 |  |
| MR | Interaction Time*Group | 3.001 | 1 | 0.083 |  | 30.5 |
|  | Group | 4.571 | 1 | **0.033** | 6.7 |  |
| CCTT1 | Interaction Time*Group | 6.989 | 1 | **0.008** | 9.5 | 14.7 |
|  | Group | 0.174 | 1 | 0.677 | 0.4 |  |
| CCTT2 | Interaction Time*Group | 8.528 | 1 | **0.003** | 14.1 | 33.6 |
|  | Group | 1.074 | 1 | 0.300 | 0.8 |  |
| Rey-1 | Interaction Time*Group | 0.473 | 1 | 0.492 |  | 23.3 |
|  | Group | 0.335 | 1 | 0.563 | 0.5 |  |
| Rey-2 | Interaction Time*Group | 0.335 | 1 | 0.563 |  | 23.3 |
|  | Group | 2.738 | 1 | 0.098 | 0.5 |  |
| Rey-3 | Interaction Time*Group | 1.690 | 1 | 0.194 |  | 23.3 |
|  | Group | 1.135 | 1 | 0.287 | 0.5 |  |
| PP-RH | Interaction Time*Group | 0.302 | 1 | 0.582 |  | 34.7 |
|  | Group | 7.002 | 1 | **0.008** | 10.0 |  |
| PP-LH | Interaction Time*Group | 0.619 | 1 | 0.431 |  | 35.8 |
|  | Group | 7.957 | 1 | **0.005** | 15.9 |  |
| PP-BH | Interaction Time*Group | 0.022 | 1 | 0.882 |  | 33.3 |
|  | Group | 8.892 | 1 | **0.003** | 16.0 |  |
| PP-Ass | Interaction Time*Group | 0.481 | 1 | 0.488 |  | 11.0 |
|  | Group | 3.994 | 1 | **0.046** | 6.9 |  |

As we were principally interested in comparing the development of the children as a function of the two musical interventions over time, we did not investigate the main effect of Time. A significant effect of time would only suggest that children of both groups showed better results at T2 than at T1, which was expected.

## 3. Results

All types of tests, measured variables, acronyms of the tests and involved abilities are resumed in Table 1. Mean descriptive data per group are provided in supplementary Table 1 (at the end of the manuscript). Raw average data can be visualized by means of boxplots of all variables and at all three time points (T0, T1 & T2) in both groups in supplementary Figure 1. The final statistical results are represented in Table 2 and illustrated in Figure 1. These final outcomes are the result of two-by-two comparisons of three different linear mixed models using likelihood-ratio tests for each comparison (see 2.9.2 Likelihood-ratio tests). Scores at T0, age, gender and handedness were controlled for by incorporating their values within the linear mixed models. The syntax of the three linear mixed models, and their comparisons by means of likelihood-ratio tests are described in the Materials and Methods section. The outcomes from the two likelihood-ratio tests respond to two questions: whether effects of 1) Interaction Time * Group and or 2) Group (OC vs Control) were significant. The interaction test responds to the question whether the two groups developed differently over time. Only estimated scores assessed by means of the linear mixed models at T1 and T2 are provided in Figure 1, as the score at T0 was incorporated in the model, in order to correct for differences at baseline between the groups, as assignment of entire classes to the OC group or Control group was random (cluster randomization).

**Figure 1.**
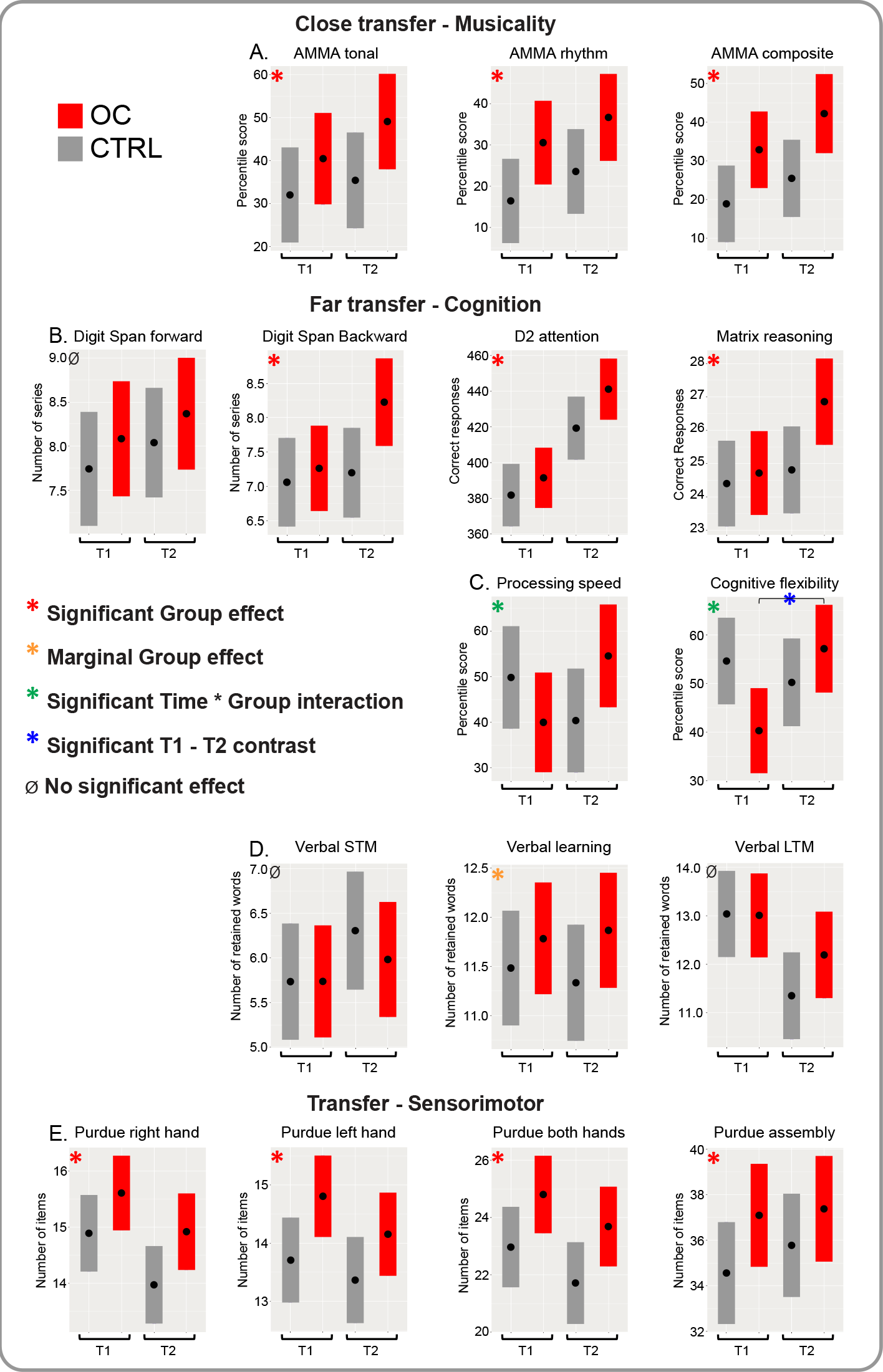
Illustration of the results provided in Table 2 for all measures. Red asterisks depict a main significant Group effect (T1 and T2 confounded), orange asterisks a marginal main Group effect (p = 0.05−0.1), green asterisks a significant Time * Group interaction, and blue asterisks a significant Bonferroni contrast OC group vs Control Group that was only computed at T2 for CCTT-1 and CCTT-2.

For the sake of brevity and transparency statistical results can be found mainly in Table 2, and as little as possible in the text. We will report on the results in detail below as a function of transfer: Near transfer (music processing), Far transfer (cognition) and Sensorimotor transfer.

### 3.1 Near transfer

All children passed the “*Advanced Measures of Music Audiation” of Gordon (AMMA;* Gordon, 1989). In the AMMA test subjects compare melodies and judge whether they are identical or that melodical/tonal or rhythmic differences occur. The OC group showed significantly enhanced percentile scores compared to the Control group, T1 and T2 confounded (from now on Group effect), for both the tonal subtest (AMMA-T) and the rhythmic subtest (AMMA-R) and thus also for the composite test (AMMA-C), that is a combination of both tonal and rhythmic scores providing an overall musicality score (see Table 2 and Figure 1 panel A). A progression in the scores can be observed between T1 and T2 (corrected for T0) in both groups. Interaction Time * Group was not significant, which means that the evolution of the scores over time did not differ significantly between the groups.

### 3.2 Far transfer

For the Digit Span Forward (DSF) and Digit Span Backward (DSB) (Wechsler, 2005) the OC group showed superior scores at T1 and T2 compared to the control group (see Table 2 and Figure 1 panel B), but the Group effect, was only significant for DSB, that reflects auditory working memory. DSF scores reflect short term auditory memory. The largest difference for DSB occurred between the groups after two full years of training.

The D2 Attention test (Brickenkamp & Zillmer, 1998), that measures selective and sustained attention and visual scanning speed (processing speed) showed a clear developmental trend for both groups, with an significant enhanced Group effect for the OC children (see Table 2 and Figure 1 panel B).

The matrix reasoning scores (MR; Wechsler, 2003) show a very similar development compared to the DSB scores, with a significant positive Group effect for the OC group, with most pronounced enhanced scores at T2.

For DSF, DSB, D2 and MR a progression in the scores can be observed between T1 and T2 in both groups. However, Interaction Time * Group was not significant, which means that the all-over evolution of the scores over time did not differ significantly between the groups.

For the Children’s Color Trails Test results are less transparent (see Table 2 and Figure 1 panel C). For both subtests (CCTT-1 measuring visual processing speed) and CCTT-2 (measuring visual processing speed and also cognitive flexibility) interaction Time * Group was significant, whereas the overall group effect did not reach significance. We can observe opposite trends in both groups: scores of the Control Group decrease from T1 to T2, but scores in the OC group increase. This is not due to baseline differences, as we controlled for T0 in the models (see supplementary Figure 1, panel C (at the end of the manuscript)).

In order to verify whether the observed progress in the scores between T1 and T2 for the OC group was significant, we applied Tukey corrected contrast between scores at T1 and T2 for CCTT-1 and CCTT-2 for both groups. For CCTT-1 no significant differences occurred in either group (OC group: t=−2.22, p= 0.219; Control Group: t=1.45, p=0.474). Tukey corrected contrasts for CCTT-2 however confirmed significant progress between T1 and T2 for the OC group (t=−3.91, p=0.012) but not for the Control group (t=0.85, p=0.83), where a marginal effect of decrease in scores occurred.

For the three subtests of the Rey Auditory Verbal Learning Test, no significant differences could be observed between the groups (see Table 2 and Figure 1 panel D). Only the second subtest (Rey-2), evaluating verbal learning showed a marginal Group effect or “trend” with higher scores for the OC group (see Table 2). Verbal short-term memory (STM, Rey-1) and Verbal long-term memory (LTM, Rey-3) did not show any significant or marginal effects. Interaction Time * Group was not significant for either of the three verbal tasks, therefore the evolution of the scores over time did not differ significantly between the groups.

### 3.3 Sensorimotor transfer

For all four subtests of the Purdue Pegboard (PP) (Lafayette, 1999) the Group effect was significant (see Table 2 and Figure 1, panel E). However, for the simple peg inserting with the right hand (PP_RH), the left hand (PP_LH) and with both hands (PP_BH), scores reached their summit already after one year of musical training in the OC group (at T1). For the more complex task, the assembly task (PP-Ass), scores increased gradually in both groups, and values were highest at T2. Interaction Time * Group was not significant for either of the four sensorimotor tasks, the evolution of the scores over time did thus not differ significantly between the groups.

## 4. Discussion

This RCT compared practicing complex instruments to sensitization to music over two full years in an intra-curricular class setting in initially non-musician children with varying backgrounds.

We could demonstrate enhanced development of different musical, cognitive and sensorimotor functions in the OC group compared to the control group. This is all the more remarkable as the courses were not individual but taught within a complete class and on four different string instruments. Moreover, the control groups received the same amount of musical education, also in a full class setting. All teachers were professional musicians. We presume that learning to master a complex instrument, as well as the ensemble playing, i.e. the dynamic interaction in the OC group, requiring the child to listen incessantly to the others and to adapt to the group and the teacher, constituted the driving force for this reinforced development.

We would like to emphasize that children would also mature and score better over time in the age groups we studied, without any musical interventions. This maturation derives partially from explicit learning within the regular school curriculum but also of spontaneous age-related acquisition of cognitive abilities in the context of natural child development (Siegler & Svetina, 2002). These superposed developmental trends may then be modulated by deliberate supplementary learning situations like musical interventions and differently so as a consequence of their nature (here focused instrumental training versus more dispersed sensitization to music).

Social economic level and other background features of the home situation of the child also play a role concerning the level of involvement in musical activities in and outside the school setting. Personality traits of both parents and the child, as well as the child’s motivation and instrument preference, may strongly influence learning and thus on subsequent transfer to other cognitive and sensorimotor achievements. (Corrigall & Schellenberg, 2015; Hannon & Trainor, 2007; Southgate & Roscigno, 2009). However, we consider that as the classes were assigned randomly to the OC and the Control programs in neighboring schools, and the study took place in public primary schools in popular neighborhoods in the Geneva area, socioeconomic and other background features of the children were balanced across the groups.

Like in the results section we will discuss effects in detail as a function of transfer.

### 4.1 Close transfer

The AMMA scores show stronger development of musical abilities that can be explained by a near transfer of learning in the OC group: intensive musical instrumental training provokes more efficient processing of musical stimuli. Potential test-retest effects were controlled for by the control group but may play a role in the all-over increasing scores between T1 and T2 in both groups (this holds for all tests). However, we reshuffled the order of the melodies at each time point, and the melodies of this task are rather abstract and not easy to memorize after a year. This test can also be used repeatedly in adult professional musicians and does thus not easily show a ceiling effect (Gordon, 2007; Gordon, 1989). For the AMMA test, auditory working memory, attention and processing speed are crucial, as two subsequent melodies have to be compared. This presumption is supported by the enhanced development of the DSB scores in the OC group. Moreover, development of visual attention and processing speed was also increased (D2 and CCTT-2 test). Assuming that visual and auditory attention and processing speed share common resources (Fougnie, Cockhren, & Marois, 2018), this then constitutes a supplementary explanation for these near transfer results. Finally, cortical and subcortical functional and structural plasticity may also explain the observed advantages for processing music in the OC group (Kraus & Strait, 2015; Strait & Kraus, 2011; Tierney et al., 2015; Trainor, Shahin, & Roberts, 2009; Zuk, Benjamin, Kenyon, & Gaab, 2014).

### 4.2 Far transfer

In addition, we observed far transfer of learning in several areas of general cognition. Far transfer of learning implies that the trained abilities go beyond the boundaries of the trained domain, although there is little consensus on the precise nature of far transfer (Barnett & Ceci, 2002). According to Barnett and Ceci, an important factor in defining far transfer is the spontaneity of its occurrence. Natural training procedures, based on real life experiences and not abstract manipulations, like musical practice or dancing, are optimal to induce generalized learning because they are complex and variable (Green & Bavelier, 2008; Green, Strobach, & Schubert, 2013) and thus have a better chance of inducing spontaneously far transfer of learning.

In the current study, this transfer to other domains may be explained by the frequent use of certain core cognitive skills that are implicated in both musical practice and other cognitive functions (Bergman Nutley et al., 2014; Roden et al., 2014), like working memory, attention and processing speed. These basic cognitive abilities are strongly involved in and thus bolstered by musical training, and then may play a role as hubs, supporting more complex cognitive abilities like matrix reasoning (Cowan, 2014). So in our point of view the transferred skills are not specific to music, but rather general (Barnett & Ceci, 2002; Thaut, 2005), they are just intensively trained during musical practice.

Working memory plays a more important role than short term memory during music practice, as one has to continuously compare what just sounded with what is coming up, and this holds for the sounds produced by the player himself as well as for the surrounding musical context.

Concerning the CCTT scores, results were somewhat controversial, with higher scores, potentially learning effects, in the control group after one year, and recovery with superior scores after two years for the OC group. Notwithstanding, development from T1 to T2 manifested for both CCTT scores exclusively in the OC group, but only reached significance for CCTT-2. It is noteworthy that the reading of musical scores plays a much more important role during the 2nd year of teaching of the OC program and may have impacted performing the CCTT-2 tasks stronger in the OC group in the second year, as score reading relies on visual scanning and processing speed, as well as cognitive flexibility. While reading the score the child has to adapt to a fluctuating auditory environment, especially in a group setting. Moreover, cognitive flexibility can be linked to enhanced sound discrimination (Saarikivi, Putkinen, Tervaniemi, & Huotilainen, 2016). In the first year the children were rather focused on holding and handling their instrument, which is very difficult in the case of string instruments, as well as on basic audio-motor processing, in the second year they can concentrate more on reading the score and listening to the others.

The absence of significant effects for the for the three subtests of the Rey Auditory Verbal Learning Test may surprise, as increase of verbal memory and other language functions is reported often as an effect of musical training (Ho, Cheung, & Chan, 2003; Jaschke et al., 2018b; Roden et al., 2012). But we should acknowledge that 1) the control group was also musically trained, and sensitization to music did show a trend for verbal IQ (Jaschke et al., 2018a), 2) children in which verbal advantages were observed started musical training at a younger age on average than the groups tested in the current experiment (Ho et al., 2003; Jaschke et al., 2018b; Roden et al., 2012) and 3) that we did find a marginal positive effect of verbal learning in the OC group as compared to the Control Group, that may have failed to reach statistical significance following lack of statistical power as our groups are of medium size.

### 4.3 Sensorimotor transfer

We would classify the sensorimotor transfer in between close and far transfer. Playing a string instrument, demanding a very asymmetric right vs left motor coordination, versus inserting or assembling small metal objects in the frontal plane, as required in our sensorimotor test (Purdue Pegboard) are not that closely related, although both require manual dexterity and bi-manual coordination.

In all four subtests the OC group outperformed the Control group. The impact of the musical training on sensorimotor performance was most obvious after the first year of training for the three simple peg inserting tasks (PP-RH, PP-LH, PP-B; see 2.4.7 Purdue Pegboard). This seems plausible, as learning to hold and handle a string instrument, involving the right and the left hand very differently is very demanding in the beginning. Especially the fine dexterity of the fingers of the left hand is particularly challenging. Concordantly, the effect size of the PP-LH and PP-BH tasks was larger than for the right hand (PP-RH; see Table 2). For the more complex task, the assembly task (PP-Ass), demanding fine finger dexterity and advanced bimanual coordination, scores increased gradually in both groups, and values were highest at T2. The control group manifested the same pattern over time, but with on average lower scores than the OC group.

### 4.4 A comparison with recent literature

Finally a recent longitudinal study over 2.5 years (Jaschke et al., 2018b) compared young primary school children (mean age 6.4 years at the beginning of the interventions) following music and arts classes one or two times per week, in addition to the regular school curriculum. Initially the study comprised three experimental groups, a music, a visual arts and a passive control group, applying cluster randomization per school. The music intervention involved acquiring basic music knowledge and listening, and also encouraged the children to play instruments, but no focused instrumental training was targeted over the full period of the intervention. Therefore, this training represents in our point of view an enriched sensitization to music. Additionally, a fourth music group was added posthoc to the study, not being part of the randomization process, consisting of children who received extra-curricular private musical instrumental lessons prior to and during the school music intervention to which they also participated. Inhibition, planning and verbal IQ improved in both music groups as compared to the art and passive control group. No differences were found between the two music groups. So, although interesting and consistent with ours, the results are to some extent contaminated by adding a posthoc group of children receiving private lessons. As the music groups were heterogeneous, with some having received music lessons prior to the study and others not, it is difficult to interpret the results or directly compare them to those of the current study.

The plus-value of the study presented here is that two groups of initially musically naïve children were compared for two different music interventions, one involving focused instrumental training and another sensitization to music with moderate practice. Our results therefore highlight the added value of intensive musical instrumental training in a group setting, encouraging general implementation in public primary schools.

## 5. Conclusion

We could show for the first time that learning to play a complex instrument in a group setting over two years, impacts positively general cognitive and sensorimotor behavior much more strongly than sensitization to music, even if the latter also comprises some musical practice. Core functions like working memory, attention, processing speed and cognitive flexibility, as well as hand dexterity and bimanual coordination, but also abstract thinking were enhanced in the Orchestra in Class group as compared to the sensitization to music group after two years of musical training.

These data show that intensive practice of a musical instrument associated with ensemble playing is a powerful means for the development of core cognitive and executive functions of the primary school child, better preparing it for secondary education. Executive functions and abstract reasoning most likely support academic achievement most strongly (Sala & Gobet, 2017). Just being sensitized to music is not sufficient to bring about such changes. The motivational and emotional aspects of musical practice could also be an explanation for the facilitation of learning (Ferreri & Verga, 2016). Making or appreciating music affects the dopaminergic and other hormone and endocrine systems (Blood & Zatorre, 2001; Chanda & Levitin, 2013; Ferreri et al., 2019). These effects are likely stronger if the music is auto-generated. Such an enriched emotional state could form a scaffold for reinforced learning.

One may wonder whether the observed benefits will remain stable over time. However, the positive influence on intelligence (IQ) of musical practice compared to other artistic activities appeared sustainable over time (Schellenberg, 2006). The authors concluded that practicing music during childhood provokes moderate but lasting positive effect on intelligence and academic performance. A recent study of twins could show that playing a musical instrument in your young years, taking into account gender, education and physical activity, reduces the risk of dementia and cognitive impairment in old age (Balbag et al., 2014), and similarly in other elderly not related individuals (Hanna-Pladdy & MacKay, 2011; White-Schwoch et al., 2013).

## 6. Limitations of the study

The age group studied here (ten to twelve years), is not ideal to show optimal benefits of musical practice and training. Neuronal plasticity is at its peak around seven years of age (Wan & Schlaug, 2010). An earlier start, but also a longer period of training could provoke stronger enhancement of development. The observed developmental enhancements in the OC group are nevertheless considerable. On the local political level of the Geneva Canton, the results generated by this study provoked a prolongation of the OC program that will now start two years earlier and last four years instead of two. Nonetheless, starting music practice during adolescence (Tierney et al., 2015) or even in elderly (Bugos, Perlstein, McCrae, Brophy, & Bedenbaugh, 2007; Dege & Kerkovius, 2018), can still provoke benefits, as our brains are plastic form the cradle to the grave.

Finally, although the control group allowed verifying for test-retest learning effects we cannot exclude that some of the progress in both groups was partially supported by such learning effects.

## 7. Acknowledgements

The authors thank Mrs. Magali Peyron, School Principal, for accepting to perform the study in her establishment and for her important contribution to the planning of the experiment. The authors are also very grateful to the children and their parents for their precious collaboration.

## 8. Author Contributions

Conceived and designed the experiment: CJ SZ MK. Passed the tests and organized the data: LA DG. Analyzed the data: CJ EDL. Wrote the paper: CJ MK SZ EDL.

## 9. Funding

This work was financed by the Accademia d’Archi – Ecole de Musique (http://www.accademia-archi.ch), with the support of CARIGEST SA in the search for anonymous sponsorship, and by the Swiss National Science Foundation (SNSF) [GZ: 100014_152841/1]. The funding sources did not play any role in study design; in the collection, analysis and interpretation of data; in the writing of the report; nor in the decision to submit the article for publication

## 10. Declaration of interest

Declarations of interest: none

---

**Supplementary Figure 1.**
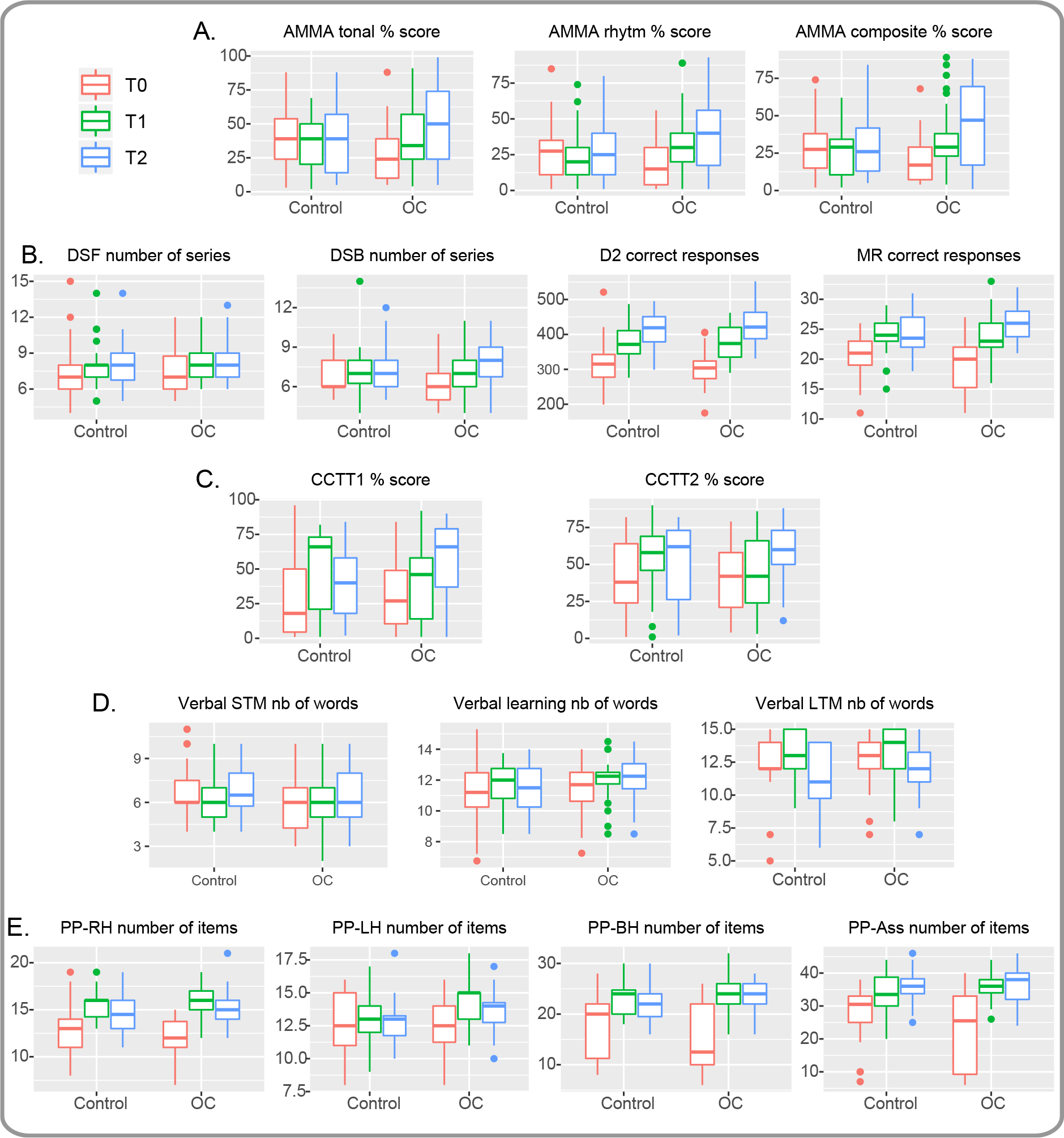
Raw data are represented as boxplots around the median, with lower and upper hinges corresponding to the first and third quartiles. The upper (respectively lower) whisker extends from the hinge to the largest (respectively smallest) value no further than 1.5 * IQR from the hinge. Outliers are represented by dots.

**Supplementary Table 1.**
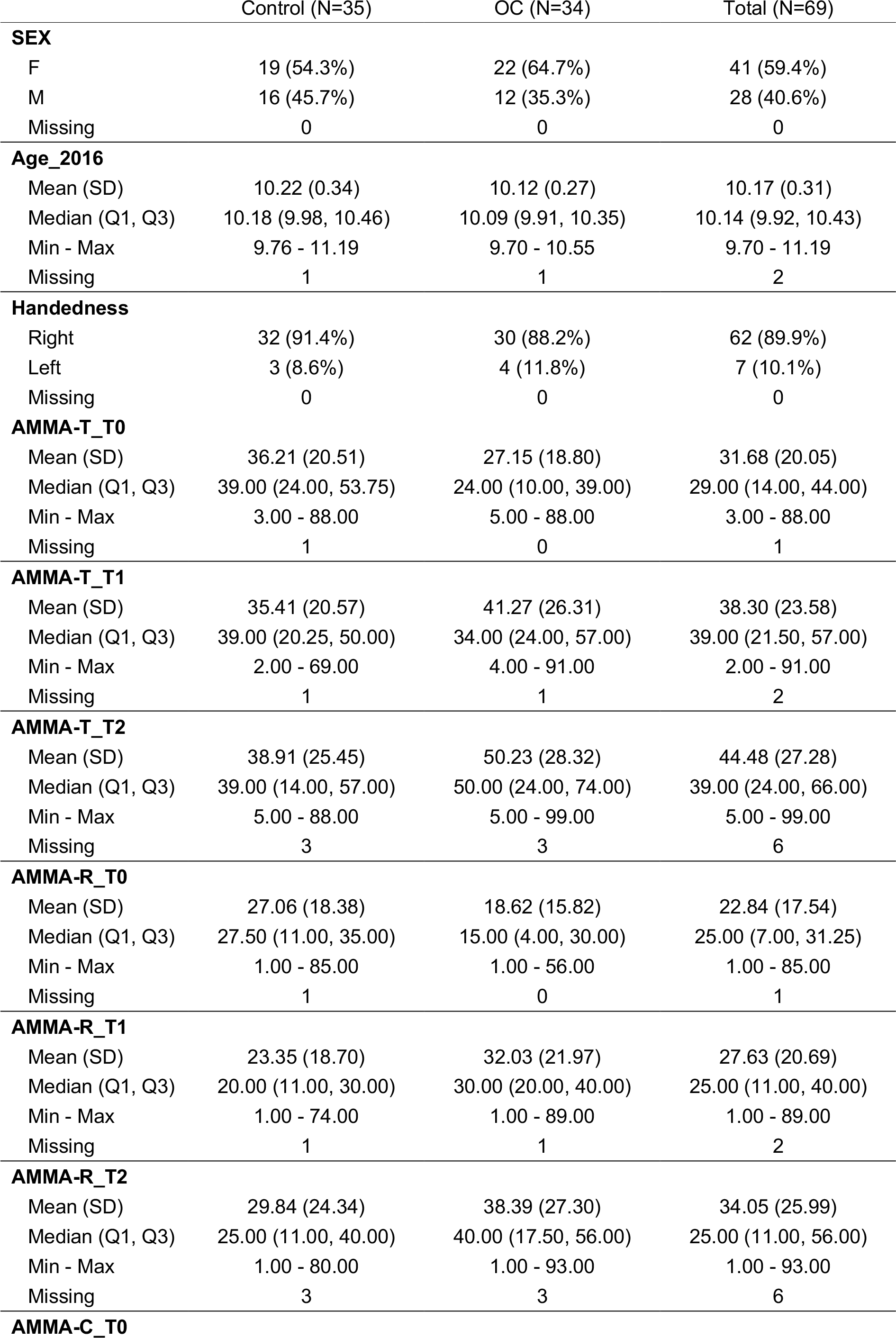
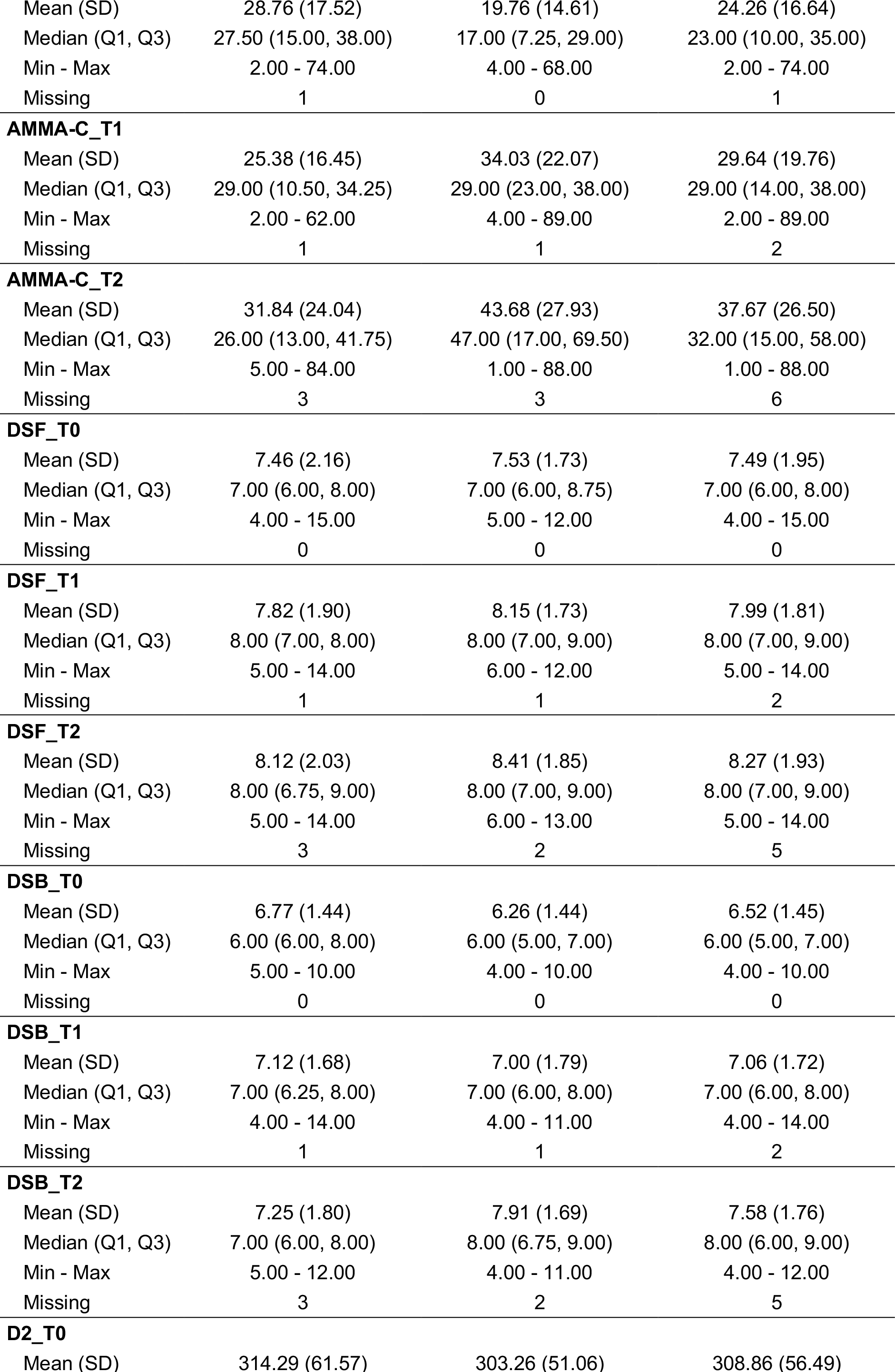
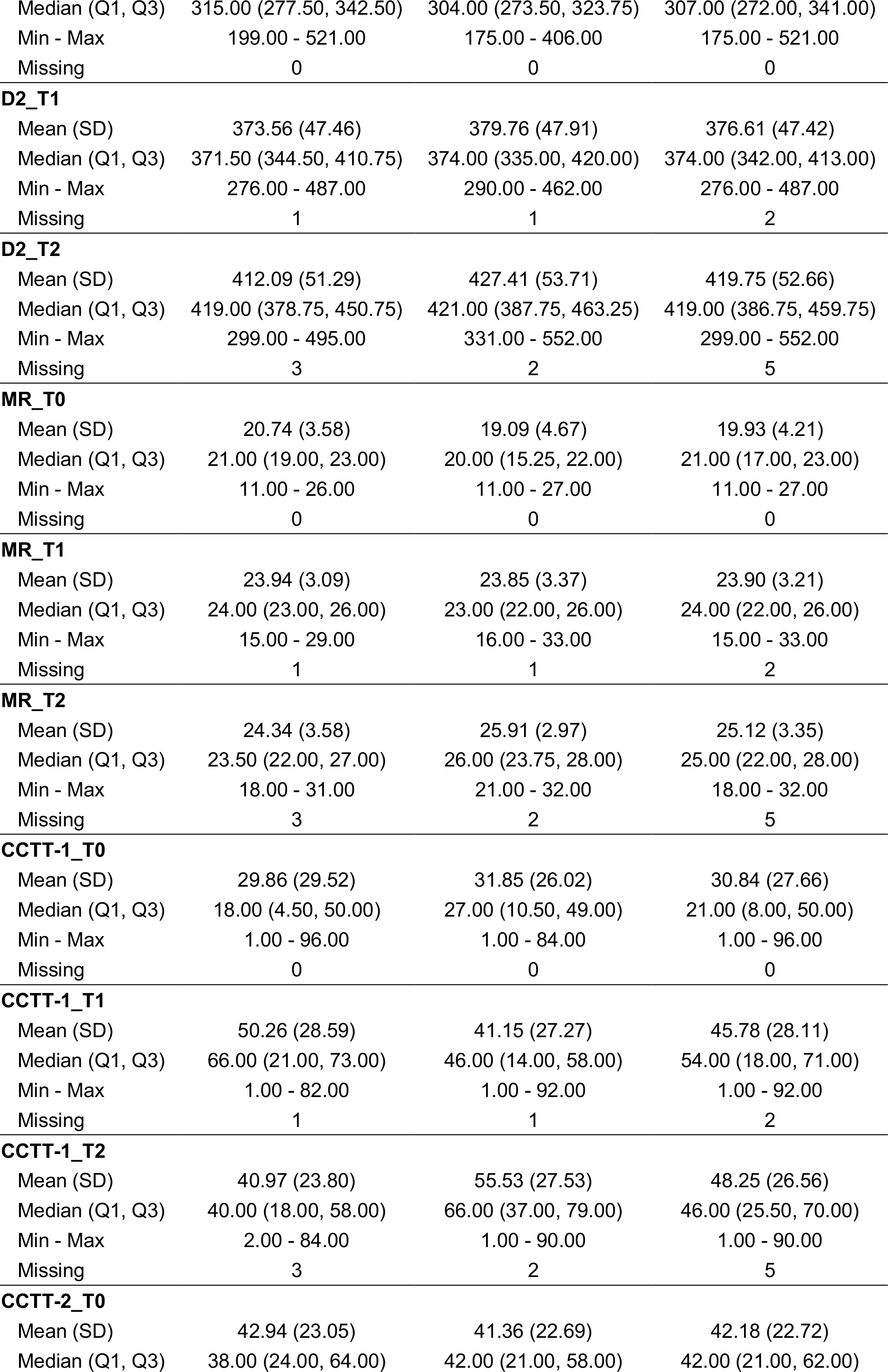
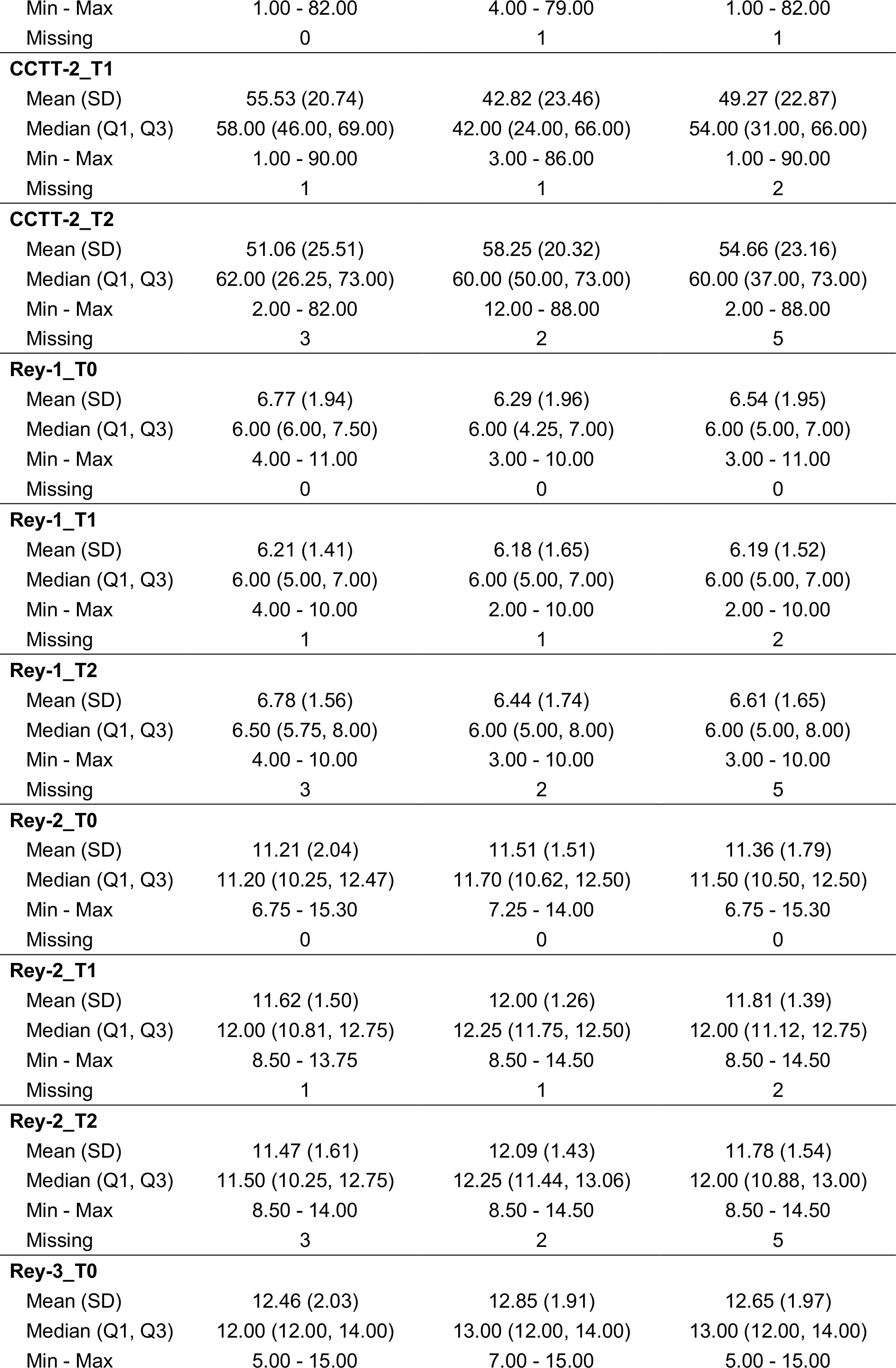
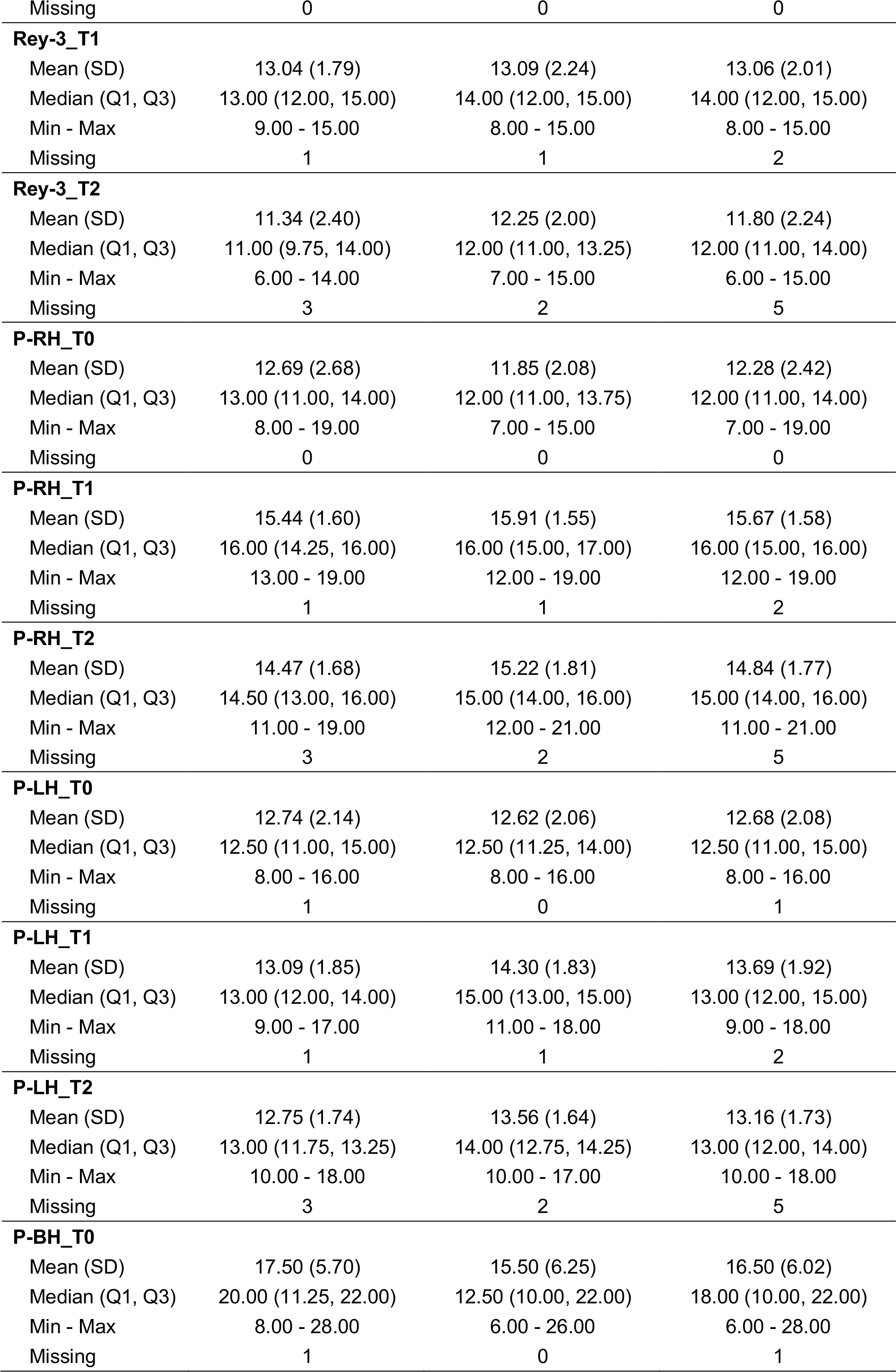
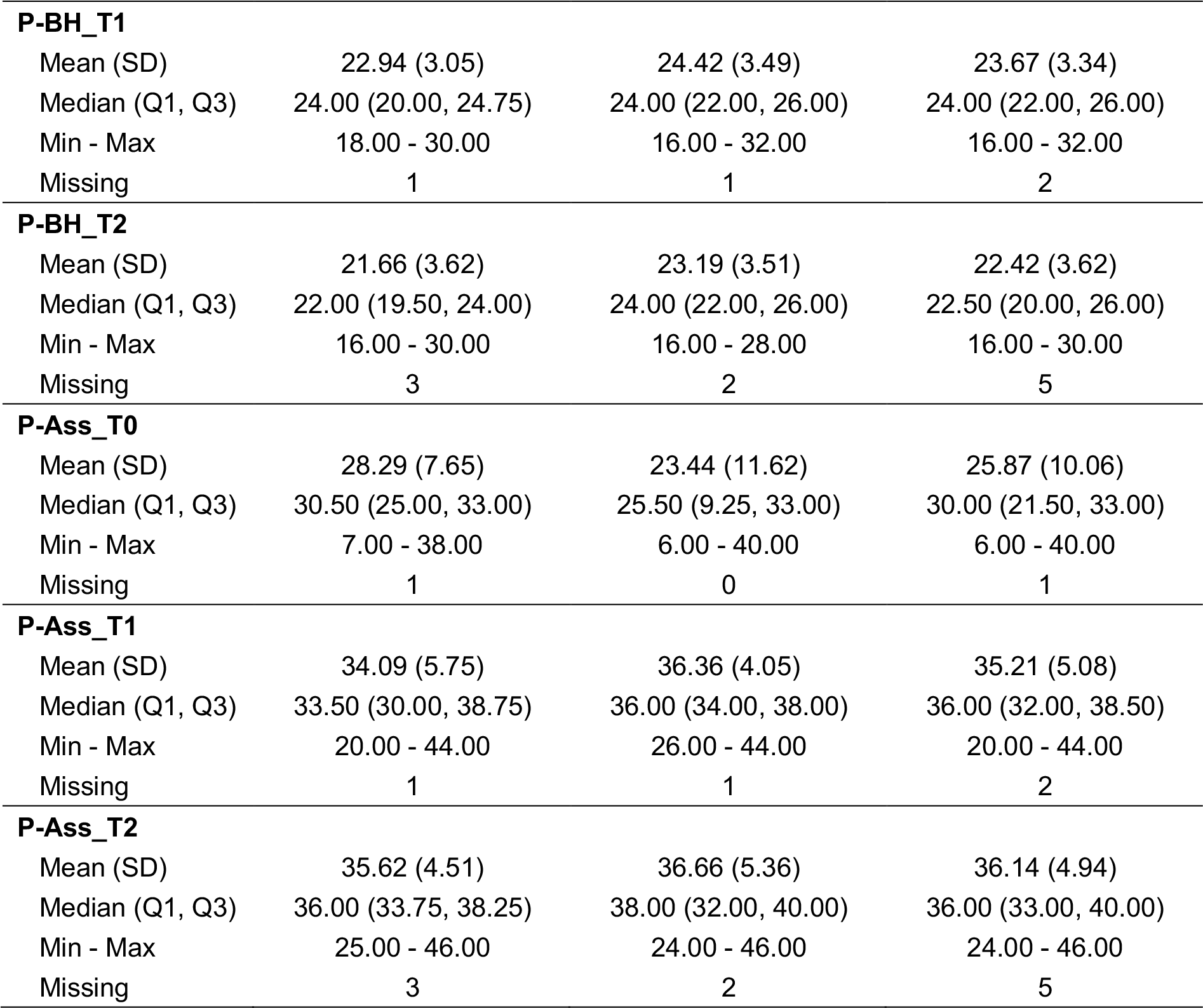
Descriptive values of all measures per group.

|  | Control (N=35) | OC (N=34) | Total (N=69) |
| --- | --- | --- | --- |
| **SEX** |  |  |  |
| F | 19 (54.3%) | 22 (64.7%) | 41 (59.4%) |
| M | 16 (45.7%) | 12 (35.3%) | 28 (40.6%) |
| Missing | 0 | 0 | 0 |
| **Age_2016** |  |  |  |
| Mean (SD) | 10.22 (0.34) | 10.12 (0.27) | 10.17 (0.31) |
| Median (Q1, Q3) | 10.18 (9.98, 10.46) | 10.09 (9.91, 10.35) | 10.14 (9.92, 10.43) |
| Min - Max | 9.76 - 11.19 | 9.70 - 10.55 | 9.70 - 11.19 |
| Missing | 1 | 1 | 2 |
| **Handedness** |  |  |  |
| Right | 32 (91.4%) | 30 (88.2%) | 62 (89.9%) |
| Left | 3 (8.6%) | 4 (11.8%) | 7 (10.1%) |
| Missing | 0 | 0 | 0 |
| **AMMA-T_T0** |  |  |  |
| Mean (SD) | 36.21 (20.51) | 27.15 (18.80) | 31.68 (20.05) |
| Median (Q1, Q3) | 39.00 (24.00, 53.75) | 24.00 (10.00, 39.00) | 29.00 (14.00, 44.00) |
| Min - Max | 3.00 - 88.00 | 5.00 - 88.00 | 3.00 - 88.00 |
| Missing | 1 | 0 | 1 |
| **AMMA-T_T1** |  |  |  |
| Mean (SD) | 35.41 (20.57) | 41.27 (26.31) | 38.30 (23.58) |
| Median (Q1, Q3) | 39.00 (20.25, 50.00) | 34.00 (24.00, 57.00) | 39.00 (21.50, 57.00) |
| Min - Max | 2.00 - 69.00 | 4.00 - 91.00 | 2.00 - 91.00 |
| Missing | 1 | 1 | 2 |
| **AMMA-T_T2** |  |  |  |
| Mean (SD) | 38.91 (25.45) | 50.23 (28.32) | 44.48 (27.28) |
| Median (Q1, Q3) | 39.00 (14.00, 57.00) | 50.00 (24.00, 74.00) | 39.00 (24.00, 66.00) |
| Min - Max | 5.00 - 88.00 | 5.00 - 99.00 | 5.00 - 99.00 |
| Missing | 3 | 3 | 6 |
| **AMMA-R_T0** |  |  |  |
| Mean (SD) | 27.06 (18.38) | 18.62 (15.82) | 22.84 (17.54) |
| Median (Q1, Q3) | 27.50 (11.00, 35.00) | 15.00 (4.00, 30.00) | 25.00 (7.00, 31.25) |
| Min - Max | 1.00 - 85.00 | 1.00 - 56.00 | 1.00 - 85.00 |
| Missing | 1 | 0 | 1 |
| **AMMA-R_T1** |  |  |  |
| Mean (SD) | 23.35 (18.70) | 32.03 (21.97) | 27.63 (20.69) |
| Median (Q1, Q3) | 20.00 (11.00, 30.00) | 30.00 (20.00, 40.00) | 25.00 (11.00, 40.00) |
| Min - Max | 1.00 - 74.00 | 1.00 - 89.00 | 1.00 - 89.00 |
| Missing | 1 | 1 | 2 |
| **AMMA-R_T2** |  |  |  |
| Mean (SD) | 29.84 (24.34) | 38.39 (27.30) | 34.05 (25.99) |
| Median (Q1, Q3) | 25.00 (11.00, 40.00) | 40.00 (17.50, 56.00) | 25.00 (11.00, 56.00) |
| Min - Max | 1.00 - 80.00 | 1.00 - 93.00 | 1.00 - 93.00 |
| Missing | 3 | 3 | 6 |
| **AMMA-C_T0** |  |  |  |
| PREPRINT |  |  |  |
| Mean (SD) | 28.76 (17.52) | 19.76 (14.61) | 24.26 (16.64) |
| Median (Q1, Q3) | 27.50 (15.00, 38.00) | 17.00 (7.25, 29.00) | 23.00 (10.00, 35.00) |
| Min - Max | 2.00 - 74.00 | 4.00 - 68.00 | 2.00 - 74.00 |
| Missing | 1 | 0 | 1 |
| **AMMA-C_T1** |  |  |  |
| Mean (SD) | 25.38 (16.45) | 34.03 (22.07) | 29.64 (19.76) |
| Median (Q1, Q3) | 29.00 (10.50, 34.25) | 29.00 (23.00, 38.00) | 29.00 (14.00, 38.00) |
| Min - Max | 2.00 - 62.00 | 4.00 - 89.00 | 2.00 - 89.00 |
| Missing | 1 | 1 | 2 |
| **AMMA-C_T2** |  |  |  |
| Mean (SD) | 31.84 (24.04) | 43.68 (27.93) | 37.67 (26.50) |
| Median (Q1, Q3) | 26.00 (13.00, 41.75) | 47.00 (17.00, 69.50) | 32.00 (15.00, 58.00) |
| Min - Max | 5.00 - 84.00 | 1.00 - 88.00 | 1.00 - 88.00 |
| Missing | 3 | 3 | 6 |
| **DSF_T0** |  |  |  |
| Mean (SD) | 7.46 (2.16) | 7.53 (1.73) | 7.49 (1.95) |
| Median (Q1, Q3) | 7.00 (6.00, 8.00) | 7.00 (6.00, 8.75) | 7.00 (6.00, 8.00) |
| Min - Max | 4.00 - 15.00 | 5.00 - 12.00 | 4.00 - 15.00 |
| Missing | 0 | 0 | 0 |
| **DSF_T1** |  |  |  |
| Mean (SD) | 7.82 (1.90) | 8.15 (1.73) | 7.99 (1.81) |
| Median (Q1, Q3) | 8.00 (7.00, 8.00) | 8.00 (7.00, 9.00) | 8.00 (7.00, 9.00) |
| Min - Max | 5.00 - 14.00 | 6.00 - 12.00 | 5.00 - 14.00 |
| Missing | 1 | 1 | 2 |
| **DSF_T2** |  |  |  |
| Mean (SD) | 8.12 (2.03) | 8.41 (1.85) | 8.27 (1.93) |
| Median (Q1, Q3) | 8.00 (6.75, 9.00) | 8.00 (7.00, 9.00) | 8.00 (7.00, 9.00) |
| Min - Max | 5.00 - 14.00 | 6.00 - 13.00 | 5.00 - 14.00 |
| Missing | 3 | 2 | 5 |
| **DSB_T0** |  |  |  |
| Mean (SD) | 6.77 (1.44) | 6.26 (1.44) | 6.52 (1.45) |
| Median (Q1, Q3) | 6.00 (6.00, 8.00) | 6.00 (5.00, 7.00) | 6.00 (5.00, 7.00) |
| Min - Max | 5.00 - 10.00 | 4.00 - 10.00 | 4.00 - 10.00 |
| Missing | 0 | 0 | 0 |
| **DSB_T1** |  |  |  |
| Mean (SD) | 7.12 (1.68) | 7.00 (1.79) | 7.06 (1.72) |
| Median (Q1, Q3) | 7.00 (6.25, 8.00) | 7.00 (6.00, 8.00) | 7.00 (6.00, 8.00) |
| Min - Max | 4.00 - 14.00 | 4.00 - 11.00 | 4.00 - 14.00 |
| Missing | 1 | 1 | 2 |
| **DSB_T2** |  |  |  |
| Mean (SD) | 7.25 (1.80) | 7.91 (1.69) | 7.58 (1.76) |
| Median (Q1, Q3) | 7.00 (6.00, 8.00) | 8.00 (6.75, 9.00) | 8.00 (6.00, 9.00) |
| Min - Max | 5.00 - 12.00 | 4.00 - 11.00 | 4.00 - 12.00 |
| Missing | 3 | 2 | 5 |
| **D2_T0** |  |  |  |
| Mean (SD) | 314.29 (61.57) | 303.26 (51.06) | 308.86 (56.49) |
| PREPRINT |  |  |  |
| Median (Q1, Q3) | 315.00 (277.50, 342.50) | 304.00 (273.50, 323.75) | 307.00 (272.00, 341.00) |
| Min - Max | 199.00 - 521.00 | 175.00 - 406.00 | 175.00 - 521.00 |
| Missing | 0 | 0 | 0 |
| **D2_T1** |  |  |  |
| Mean (SD) | 373.56 (47.46) | 379.76 (47.91) | 376.61 (47.42) |
| Median (Q1, Q3) | 371.50 (344.50, 410.75) | 374.00 (335.00, 420.00) | 374.00 (342.00, 413.00) |
| Min - Max | 276.00 - 487.00 | 290.00 - 462.00 | 276.00 - 487.00 |
| Missing | 1 | 1 | 2 |
| **D2_T2** |  |  |  |
| Mean (SD) | 412.09 (51.29) | 427.41 (53.71) | 419.75 (52.66) |
| Median (Q1, Q3) | 419.00 (378.75, 450.75) | 421.00 (387.75, 463.25) | 419.00 (386.75, 459.75) |
| Min - Max | 299.00 - 495.00 | 331.00 - 552.00 | 299.00 - 552.00 |
| Missing | 3 | 2 | 5 |
| **MR_T0** |  |  |  |
| Mean (SD) | 20.74 (3.58) | 19.09 (4.67) | 19.93 (4.21) |
| Median (Q1, Q3) | 21.00 (19.00, 23.00) | 20.00 (15.25, 22.00) | 21.00 (17.00, 23.00) |
| Min - Max | 11.00 - 26.00 | 11.00 - 27.00 | 11.00 - 27.00 |
| Missing | 0 | 0 | 0 |
| **MR_T1** |  |  |  |
| Mean (SD) | 23.94 (3.09) | 23.85 (3.37) | 23.90 (3.21) |
| Median (Q1, Q3) | 24.00 (23.00, 26.00) | 23.00 (22.00, 26.00) | 24.00 (22.00, 26.00) |
| Min - Max | 15.00 - 29.00 | 16.00 - 33.00 | 15.00 - 33.00 |
| Missing | 1 | 1 | 2 |
| **MR_T2** |  |  |  |
| Mean (SD) | 24.34 (3.58) | 25.91 (2.97) | 25.12 (3.35) |
| Median (Q1, Q3) | 23.50 (22.00, 27.00) | 26.00 (23.75, 28.00) | 25.00 (22.00, 28.00) |
| Min - Max | 18.00 - 31.00 | 21.00 - 32.00 | 18.00 - 32.00 |
| Missing | 3 | 2 | 5 |
| **CCTT-1_T0** |  |  |  |
| Mean (SD) | 29.86 (29.52) | 31.85 (26.02) | 30.84 (27.66) |
| Median (Q1, Q3) | 18.00 (4.50, 50.00) | 27.00 (10.50, 49.00) | 21.00 (8.00, 50.00) |
| Min - Max | 1.00 - 96.00 | 1.00 - 84.00 | 1.00 - 96.00 |
| Missing | 0 | 0 | 0 |
| **CCTT-1_T1** |  |  |  |
| Mean (SD) | 50.26 (28.59) | 41.15 (27.27) | 45.78 (28.11) |
| Median (Q1, Q3) | 66.00 (21.00, 73.00) | 46.00 (14.00, 58.00) | 54.00 (18.00, 71.00) |
| Min - Max | 1.00 - 82.00 | 1.00 - 92.00 | 1.00 - 92.00 |
| Missing | 1 | 1 | 2 |
| **CCTT-1_T2** |  |  |  |
| Mean (SD) | 40.97 (23.80) | 55.53 (27.53) | 48.25 (26.56) |
| Median (Q1, Q3) | 40.00 (18.00, 58.00) | 66.00 (37.00, 79.00) | 46.00 (25.50, 70.00) |
| Min - Max | 2.00 - 84.00 | 1.00 - 90.00 | 1.00 - 90.00 |
| Missing | 3 | 2 | 5 |
| **CCTT-2_T0** |  |  |  |
| Mean (SD) | 42.94 (23.05) | 41.36 (22.69) | 42.18 (22.72) |
| Median (Q1, Q3) | 38.00 (24.00, 64.00) | 42.00 (21.00, 58.00) | 42.00 (21.00, 62.00) |
| PREPRINT |  |  |  |
| Min - Max | 1.00 - 82.00 | 4.00 - 79.00 | 1.00 - 82.00 |
| Missing | 0 | 1 | 1 |
| **CCTT-2_T1** |  |  |  |
| Mean (SD) | 55.53 (20.74) | 42.82 (23.46) | 49.27 (22.87) |
| Median (Q1, Q3) | 58.00 (46.00, 69.00) | 42.00 (24.00, 66.00) | 54.00 (31.00, 66.00) |
| Min - Max | 1.00 - 90.00 | 3.00 - 86.00 | 1.00 - 90.00 |
| Missing | 1 | 1 | 2 |
| **CCTT-2_T2** |  |  |  |
| Mean (SD) | 51.06 (25.51) | 58.25 (20.32) | 54.66 (23.16) |
| Median (Q1, Q3) | 62.00 (26.25, 73.00) | 60.00 (50.00, 73.00) | 60.00 (37.00, 73.00) |
| Min - Max | 2.00 - 82.00 | 12.00 - 88.00 | 2.00 - 88.00 |
| Missing | 3 | 2 | 5 |
| **Rey-1_T0** |  |  |  |
| Mean (SD) | 6.77 (1.94) | 6.29 (1.96) | 6.54 (1.95) |
| Median (Q1, Q3) | 6.00 (6.00, 7.50) | 6.00 (4.25, 7.00) | 6.00 (5.00, 7.00) |
| Min - Max | 4.00 - 11.00 | 3.00 - 10.00 | 3.00 - 11.00 |
| Missing | 0 | 0 | 0 |
| **Rey-1_T1** |  |  |  |
| Mean (SD) | 6.21 (1.41) | 6.18 (1.65) | 6.19 (1.52) |
| Median (Q1, Q3) | 6.00 (5.00, 7.00) | 6.00 (5.00, 7.00) | 6.00 (5.00, 7.00) |
| Min - Max | 4.00 - 10.00 | 2.00 - 10.00 | 2.00 - 10.00 |
| Missing | 1 | 1 | 2 |
| **Rey-1_T2** |  |  |  |
| Mean (SD) | 6.78 (1.56) | 6.44 (1.74) | 6.61 (1.65) |
| Median (Q1, Q3) | 6.50 (5.75, 8.00) | 6.00 (5.00, 8.00) | 6.00 (5.00, 8.00) |
| Min - Max | 4.00 - 10.00 | 3.00 - 10.00 | 3.00 - 10.00 |
| Missing | 3 | 2 | 5 |
| **Rey-2_T0** |  |  |  |
| Mean (SD) | 11.21 (2.04) | 11.51 (1.51) | 11.36 (1.79) |
| Median (Q1, Q3) | 11.20 (10.25, 12.47) | 11.70 (10.62, 12.50) | 11.50 (10.50, 12.50) |
| Min - Max | 6.75 - 15.30 | 7.25 - 14.00 | 6.75 - 15.30 |
| Missing | 0 | 0 | 0 |
| **Rey-2_T1** |  |  |  |
| Mean (SD) | 11.62 (1.50) | 12.00 (1.26) | 11.81 (1.39) |
| Median (Q1, Q3) | 12.00 (10.81, 12.75) | 12.25 (11.75, 12.50) | 12.00 (11.12, 12.75) |
| Min - Max | 8.50 - 13.75 | 8.50 - 14.50 | 8.50 - 14.50 |
| Missing | 1 | 1 | 2 |
| **Rey-2_T2** |  |  |  |
| Mean (SD) | 11.47 (1.61) | 12.09 (1.43) | 11.78 (1.54) |
| Median (Q1, Q3) | 11.50 (10.25, 12.75) | 12.25 (11.44, 13.06) | 12.00 (10.88, 13.00) |
| Min - Max | 8.50 - 14.00 | 8.50 - 14.50 | 8.50 - 14.50 |
| Missing | 3 | 2 | 5 |
| **Rey-3_T0** |  |  |  |
| Mean (SD) | 12.46 (2.03) | 12.85 (1.91) | 12.65 (1.97) |
| Median (Q1, Q3) | 12.00 (12.00, 14.00) | 13.00 (12.00, 14.00) | 13.00 (12.00, 14.00) |
| Min - Max | 5.00 - 15.00 | 7.00 - 15.00 | 5.00 - 15.00 |

**Supplementary Table 1**
|  |  |  |  |
| --- | --- | --- | --- |
| Missing | 0 | 0 | 0 |
| **Rey-3_T1** |  |  |  |
| Mean (SD) | 13.04 (1.79) | 13.09 (2.24) | 13.06 (2.01) |
| Median (Q1, Q3) | 13.00 (12.00, 15.00) | 14.00 (12.00, 15.00) | 14.00 (12.00, 15.00) |
| Min - Max | 9.00 - 15.00 | 8.00 - 15.00 | 8.00 - 15.00 |
| Missing | 1 | 1 | 2 |
| **Rey-3_T2** |  |  |  |
| Mean (SD) | 11.34 (2.40) | 12.25 (2.00) | 11.80 (2.24) |
| Median (Q1, Q3) | 11.00 (9.75, 14.00) | 12.00 (11.00, 13.25) | 12.00 (11.00, 14.00) |
| Min - Max | 6.00 - 14.00 | 7.00 - 15.00 | 6.00 - 15.00 |
| Missing | 3 | 2 | 5 |
| **P-RH_T0** |  |  |  |
| Mean (SD) | 12.69 (2.68) | 11.85 (2.08) | 12.28 (2.42) |
| Median (Q1, Q3) | 13.00 (11.00, 14.00) | 12.00 (11.00, 13.75) | 12.00 (11.00, 14.00) |
| Min - Max | 8.00 - 19.00 | 7.00 - 15.00 | 7.00 - 19.00 |
| Missing | 0 | 0 | 0 |
| **P-RH_T1** |  |  |  |
| Mean (SD) | 15.44 (1.60) | 15.91 (1.55) | 15.67 (1.58) |
| Median (Q1, Q3) | 16.00 (14.25, 16.00) | 16.00 (15.00, 17.00) | 16.00 (15.00, 16.00) |
| Min - Max | 13.00 - 19.00 | 12.00 - 19.00 | 12.00 - 19.00 |
| Missing | 1 | 1 | 2 |
| **P-RH_T2** |  |  |  |
| Mean (SD) | 14.47 (1.68) | 15.22 (1.81) | 14.84 (1.77) |
| Median (Q1, Q3) | 14.50 (13.00, 16.00) | 15.00 (14.00, 16.00) | 15.00 (14.00, 16.00) |
| Min - Max | 11.00 - 19.00 | 12.00 - 21.00 | 11.00 - 21.00 |
| Missing | 3 | 2 | 5 |
| **P-LH_T0** |  |  |  |
| Mean (SD) | 12.74 (2.14) | 12.62 (2.06) | 12.68 (2.08) |
| Median (Q1, Q3) | 12.50 (11.00, 15.00) | 12.50 (11.25, 14.00) | 12.50 (11.00, 15.00) |
| Min - Max | 8.00 - 16.00 | 8.00 - 16.00 | 8.00 - 16.00 |
| Missing | 1 | 0 | 1 |
| **P-LH_T1** |  |  |  |
| Mean (SD) | 13.09 (1.85) | 14.30 (1.83) | 13.69 (1.92) |
| Median (Q1, Q3) | 13.00 (12.00, 14.00) | 15.00 (13.00, 15.00) | 13.00 (12.00, 15.00) |
| Min - Max | 9.00 - 17.00 | 11.00 - 18.00 | 9.00 - 18.00 |
| Missing | 1 | 1 | 2 |
| **P-LH_T2** |  |  |  |
| Mean (SD) | 12.75 (1.74) | 13.56 (1.64) | 13.16 (1.73) |
| Median (Q1, Q3) | 13.00 (11.75, 13.25) | 14.00 (12.75, 14.25) | 13.00 (12.00, 14.00) |
| Min - Max | 10.00 - 18.00 | 10.00 - 17.00 | 10.00 - 18.00 |
| Missing | 3 | 2 | 5 |
| **P-BH_T0** |  |  |  |
| Mean (SD) | 17.50 (5.70) | 15.50 (6.25) | 16.50 (6.02) |
| Median (Q1, Q3) | 20.00 (11.25, 22.00) | 12.50 (10.00, 22.00) | 18.00 (10.00, 22.00) |
| Min - Max | 8.00 - 28.00 | 6.00 - 26.00 | 6.00 - 28.00 |
| Missing | 1 | 0 | 1 |
| **P-BH_T1** |  |  |  |
| Mean (SD) | 22.94 (3.05) | 24.42 (3.49) | 23.67 (3.34) |
| Median (Q1, Q3) | 24.00 (20.00, 24.75) | 24.00 (22.00, 26.00) | 24.00 (22.00, 26.00) |
| Min - Max | 18.00 - 30.00 | 16.00 - 32.00 | 16.00 - 32.00 |
| Missing | 1 | 1 | 2 |
| **P-BH_T2** |  |  |  |
| Mean (SD) | 21.66 (3.62) | 23.19 (3.51) | 22.42 (3.62) |
| Median (Q1, Q3) | 22.00 (19.50, 24.00) | 24.00 (22.00, 26.00) | 22.50 (20.00, 26.00) |
| Min - Max | 16.00 - 30.00 | 16.00 - 28.00 | 16.00 - 30.00 |
| Missing | 3 | 2 | 5 |
| **P-Ass_T0** |  |  |  |
| Mean (SD) | 28.29 (7.65) | 23.44 (11.62) | 25.87 (10.06) |
| Median (Q1, Q3) | 30.50 (25.00, 33.00) | 25.50 (9.25, 33.00) | 30.00 (21.50, 33.00) |
| Min - Max | 7.00 - 38.00 | 6.00 - 40.00 | 6.00 - 40.00 |
| Missing | 1 | 0 | 1 |
| **P-Ass_T1** |  |  |  |
| Mean (SD) | 34.09 (5.75) | 36.36 (4.05) | 35.21 (5.08) |
| Median (Q1, Q3) | 33.50 (30.00, 38.75) | 36.00 (34.00, 38.00) | 36.00 (32.00, 38.50) |
| Min - Max | 20.00 - 44.00 | 26.00 - 44.00 | 20.00 - 44.00 |
| Missing | 1 | 1 | 2 |
| **P-Ass_T2** |  |  |  |
| Mean (SD) | 35.62 (4.51) | 36.66 (5.36) | 36.14 (4.94) |
| Median (Q1, Q3) | 36.00 (33.75, 38.25) | 38.00 (32.00, 40.00) | 36.00 (33.00, 40.00) |
| Min - Max | 25.00 - 46.00 | 24.00 - 46.00 | 24.00 - 46.00 |
| Missing | 3 | 2 | 5 |

## Notes

https://yareta.unige.ch

